# Coupling of renal sodium and calcium transport: A modeling analysis

**DOI:** 10.1101/2023.05.29.542749

**Authors:** Shervin Hakimi, Pritha Dutta, Anita T. Layton

## Abstract

Calcium (Ca^2+^) transport along the nephron occurs via specific transcellular and paracellular pathways, and is coupled to the transport of other electrolytes. Notably sodium (Na^+^) transport establishes an electrochemical gradient to drive Ca^2+^ reabsorption. Hence, alterations in renal Na^+^ handling, under pathophysiological conditions or pharmacological manipulations, can have major effects on Ca^2+^ transport. An important class of pharmacological agent is diuretics, which are commonly prescribed for the management of blood pressure and fluid balance. The pharmacological targets of diuretics generally directly facilitate Na^+^ transport, but also indirectly affect renal Ca^2+^ handling. To better understand the underlying mechanisms, we have developed a computational model of electrolyte transport along the superficial nephron in the kidney of a male and female rat. Sex differences in renal Ca^2+^ handling are represented. Model simulations predict in the female rat nephron lower Ca^2+^ reabsorption in the proximal tubule and thick ascending limb, but higher reabsorption in the late distal convoluted tubule and connecting tubule, compared to the male nephron. The male rat kidney model yields a higher urinary calcium excretion than female, consistent with animal experiments. Additionally, we conducted simulations of inhibition of channels and transporters that play a major role in Na^+^ and Ca^2+^ transport. Model results indicate that along the proximal tubule and ascending thick ascending limb, Ca^2+^ and Na^+^ transport occurs in parallel, but those processes are dissociated in the distal convoluted tubule. Simulations also reveal sex-specific responses to different pharmacological manipulations.

## Introduction

Calcium is essential for many physiological and cellular processes, including muscle contraction, blood clotting, enzyme activation, cell differentiation, immune response, programmed cell death, and neuronal activity.^1^ The kidney plays an important role in the maintenance of calcium balance in the body by regulating calcium reabsorption and excretion. Calcium ions are filtered by the glomeruli, and <2% are excreted in urine.^2^ The majority of Ca^2+^ reabsorption along the renal tubule occurs via a passive paracellular Ca^2+^ transport process in the proximal tubule through claudin-2 and -12 pores (Cldn2, Cldn12) and the thick ascending limb of the loop of Henle via claudin-16 and -19 pores (Cldn16 and Cldn19).^3^ Approximately 10% of Ca^2+^ is reabsorbed via a transcellular Ca^2+^ transport mechanism in the distal part of the nephron, with apical entry mediated by transient receptor potential vanilloid subtype V (TRVP5).^2, 3^ Paracellular Ca^2+^ transport is driven by the transepithelial electrochemical gradient that is generated by Na^+^ and water reabsorption and is only indirectly regulated by hormones.^2^ In contrast, the transcellular pathway is specifically controlled by calciotropic hormones, including parathyroid hormone (PTH), Fibroblast growth factor 23 (FGF23), and 1,25-dihydroxy vitamin D_3_ [1,25(OH)_2_D_3_].^2, 4^ Due to this regulatory process, an organism can respond to fluctuations in dietary Ca^2+^ and adapt to changes in demand during certain physiological states, such as growth, pregnancy, lactation, and aging. Some of these physiological states are unique to females.

Sex differences have also been observed in Ca^2+^ excretion after an oral challenge, and in the association of urinary Ca^2+^ excretion with serum Ca^2+^ and vitamin D levels.^5, 6^ Given the essential role of the kidney in Ca^2+^ homeostasis, sex differences in Ca^2+^ response may be attributable, in part, to differences in kidney structure and function. Here, we focused on sex differences in renal electrolyte transporters, channels, and claudins (which we collectively refer to as transporters). Veiras et al.^7^ reported sexual dimorphism in transporter abundance patterns in rodents. Their findings revealed markedly different transport capacity in the proximal tubule of male and female rat nephrons. Compared with male rat nephrons, female rat nephrons exhibit, in the proximal tubule, greater Na^+^/H^+^ exchanger 3 (NHE3) phosphorylation and redistribution to the base of the microvilli, where activity is lower,^8^ as well as lower abundance of Na^+^-P_i_ cotransporter 2 (NaPi2), Aquaporin 1 (AQP1), and Cldn2 . Consequently, the proximal tubule in the female rat reabsorbs a substantially lower fraction of filtered Na^+^, compared to the male rat.^7^ Downstream of the macula densa, female rats exhibit higher abundance and phosphorylation of NCC and more abundant ENaC and claudin-7.^7^

As noted above, paracellular Ca^2+^ transport is driven by the transepithelial electrochemical gradient generated by Na^+^ and water reabsorption. Thus, alterations in tubular Na^+^ transport can have profound effects on renal Ca^2+^ handling. With the exception of the thin descending limb of the loop of Henle, all renal tubular segments participate significantly in the retrieval of filtered Na^+^. Consequently, any impairment in Na^+^ transport can alter urinary Na^+^ and Ca^2+^ excretion, and therefore affect arterial blood pressure and Ca^2+^ homeostasis. NHE3 is the major apical Na^+^ transporter in the proximal tubule,^9, 10^ which is responsible for the majority of renal Na^+^ reabsorption,^11^ and contributes to Na^+^ reabsorption in the thick ascending limb. The pacemaker of Na^+^ transport in the thick ascending limb, however, is the bumetanide-sensitive Na^+^-K^+^-2Cl^−^ cotransporter (NKCC2).^12^ The importance of NKCC2 for Na^+^ balance is evinced by the extensive use of loop diuretics in the treatment of acute edematous states and hypertension.^12^ Additionally, inactivating mutations in the NKCC2 gene *SLC12A1* cause type I Bartter syndrome, a life-threatening disease featuring arterial hypotension along with electrolyte balance abnormalities.^12^ Na^+^ uptake across the apical membrane of distal convoluted tubules and collecting duct principal cells is mediated mainly by NCC and ENaC, respectively.^13^ The importance of NCC and ENaC is underlined by the common use of their pharmacologic inhibitors, thiazide diuretics and potassium-sparing diuretics such as amiloride, respectively, to treat hypertension.^14^

In addition to the pharmacological inhibition of renal Na^+^ reabsorption by several specific drugs such as diuretics, the activity of these transporters may also be inhibited by several endogenous agents such as dopamine, endothelin, parathyroid hormone, adenosine, and adenosine triphosphate (ATP).^15–20^ Generally, the pharmacological targets of diuretics directly facilitate Na^+^ transport, but they may also indirectly affect renal Ca^2+^ handling, i.e., by establishing a prerequisite electrochemical gradient. It is therefore not surprising that substantial alterations in Ca^2+^ handling can be observed following diuretic treatment.^21^ Thus, a goal of this study is to investigate the extent to which inhibitors of Na^+^ transporter along the nephron alter transepithelial Ca^2+^ transport, how these effects vary among different nephron segments, and how they differ between the sexes.

## Methods

We previously developed an epithelial cell-based model of solute transport along the superficial nephron in the kidney of a male and female rat. The model nephron extends from Bowman’s capsule to the papillary tip. Baseline single nephron glomerular filtration rate (SNGFR) is set to 30 and 24 nL/min in male and female rats, respectively. We expanded the model by adding Ca^2+^ to the solutes represented, so that the model accounts for 16 solutes: Na^+^, Ca^2+^, K^+^, Cl^−^, HCO_3_^−^, H_2_CO_3_, CO_2_, NH_3_, NH_4_^+^, HPO_4_^2−^, H_2_PO_4_^−^, H^+^, HCO_2_^−^, H_2_CO_2_, urea, and glucose. The model is formulated for the steady state and predicts luminal fluid flow, hydrostatic pressure, luminal fluid solute concentrations, and, except for the descending limb segment, cytosolic solute concentrations, membrane potential, and transcellular and paracellular fluxes. A schematic diagram for the model is shown in Fig. 1.

**Figure 1.**
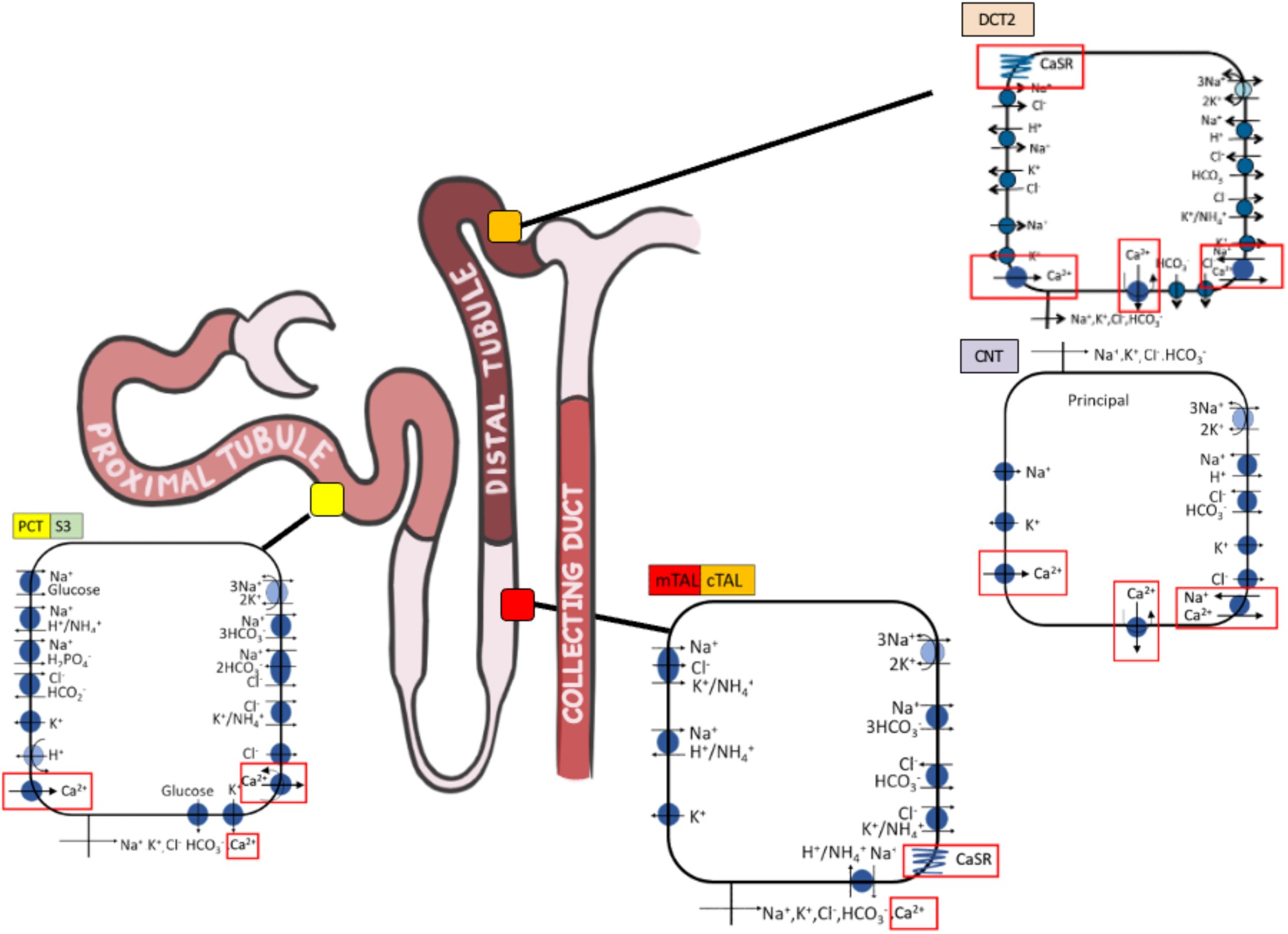
Illustration of a nephron and cells where Ca^2+^ reabsorption primarily occurs. Ca^2+^ membrane transporters and paracellular fluxes are highlighted in red boxes. PCT, proximal convoluted tubule; mTAL, medullary thick ascending limb; cTAL, cortical thick ascending limb; DCT, distal convoluted tubule; CNT, connecting tubule. DCT2 cell and the principal cell of the CNT tubule are represented (Ca^2+^-specific channels and transporters in DCT2 are similar to CNT).

*Simulating transient receptor potential vanilloid subtype V (TRPV5)-mediated Ca^2+^ transport.* The late distal tubule cells and the principal cells of the connecting tubule reabsorb Ca^2+^ entirely via the transcellular pathway, facilitated by the TRPV5 protein located at the apical membrane of these cells. The TRPV5-mediated Ca^2+^ flux is computed using the modified form of Ohm’s law:

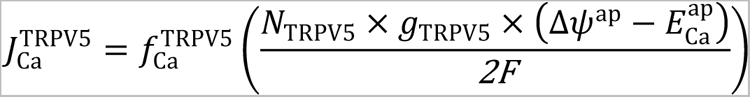

*N*_TRPV5_ is the TRPV5 channel density, *g*_TRPV5_ is the channel conductance whose value depends on the intracellular pH (pH_i_) and extracellular pH (pH_e_) ^22^, Δ*ψ*^*ap*^ is the electrical potential difference across the apical membrane (lumen minus cell), and 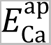 is the Nernst potential of Ca^2+^, given by

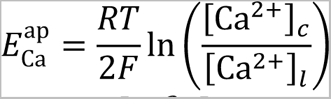

where the subscripts *c* and *l* associated with [Ca^2+^] denote the cellular and luminal compartments, respectively. 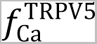 captures the Ca^2+^-induced inhibition of TRPV5 activity, given by

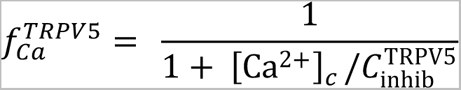

where 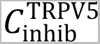 denotes the half-maximal value for [Ca^2+^]*_c_*

Cellular calcium extrusion involves two key proteins located at the basolateral membranes: Type 1 Sodium-Calcium Exchanger (NCX1) and plasma membrane Ca^2+^-ATPase (PMCA). PMCA is present in proximal tubule, late distal tubule, and connecting tubule cells, while NCX1 is exclusive to late distal tubule and connecting tubule cells.^23, 24^ In these cells, NCX1 accounts for around 70% of extrusion, with PMCA contributing 30%.^25^

### Simulating NCX1 mediated Ca^2+^ transport

The NCX1.3 isoform is the predominant isoform of NCX1 in DCT2-CNT cells. However, due to the absence of existing kinetic models for NCX1.3, the current model utilizes the NCX1.1 isoform, which is the predominant isoform in cardiac cells. The NCX1-mediated fluxes are given by:

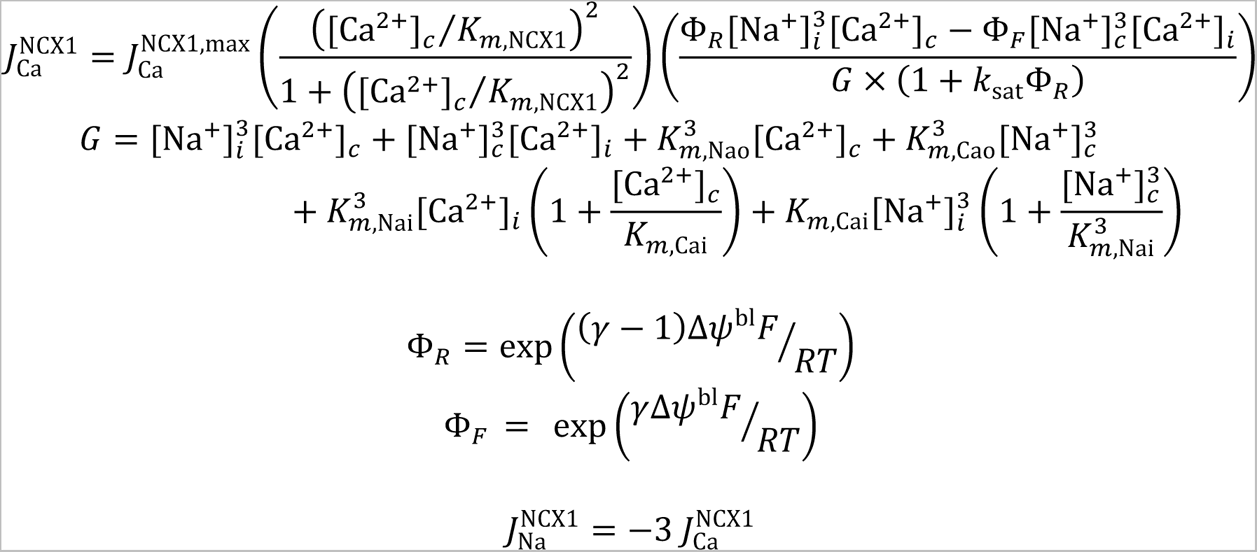

where parameters in the above equations are described in Table 3.

### Simulating PMCA mediated Ca^2+^ transport

The PMCA-mediated Ca^2+^ flux 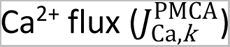 for segment *k* is given by

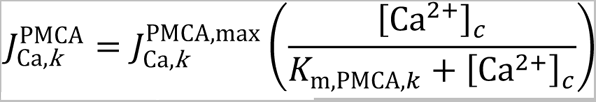

where 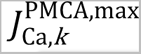 represents the maximum PMCA flux, *K*_m,PMCA,*k*_ denotes the affinity of the pump to Ca^2+^.

### Simulating transport regulation by Ca^2+^-sensing receptor (CaSR)

Expressed by the thick ascending limb, distal convoluted tubule, and medullary collecting duct cells, CaSR regulates a number of transport processes. Its action is modelled by the scaling factor ^26^

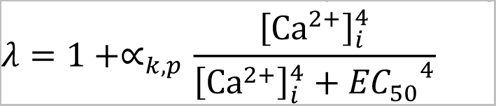

where *EC*_50_ denotes the maximum half concentration and is equal to 1.25 mM,^27^ and [Ca^+,^]_C_ represents Ca^2+^ concentration in compartment *i*. α_*k,p*_ is a scaling coefficient for the effect associated with segment *k* and transporter *p*, with positive and negative sign representing activation and inhibition, respectively.

Expressed on the basolateral membrane of the thick ascending limb cells, CaSR modulates tight-junction Ca^2+^ permeability in response to changes in serum [Ca^2+^]; thus, the subscript *i = s*. To represent the inhibitory effect, 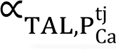 is taken to be -0.5714. The tight-junction Ca^2+^ permeability is given by 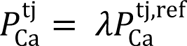, where 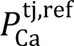 denotes 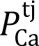 at zero serum [Ca^2+^]. CaSR also affects the activity of non-calcium-specific channels and transporters: *In vivo* experiments in rats have shown that increasing serum [Ca^2+^] from 1.1 to 5 mM reduces renal outer medullary potassium channel (ROMK) activity by 84%. In *in vivo* experiments in male mice, calcimimetic treatment reduces NKCC2 phosphorylation by ∼50%, without changing NKCC2 expression levels^28^. Given these observations, we set α_TAL,ROMK_ to -0.84 and α_TAL,NKCC2_ to -0.5.

CaSR is also expressed at the apical membrane of distal convoluted tubule cells; thus, *i = l* indicates that CaSR responds to luminal [Ca^2+^]. *In vivo* experiments on male mice have revealed the activatory effect of CaSR on NCC. Thus, α_DCT,NCC_ is taken to be 0.5, with the positive sign indicating activatory action. Further downstream, CaSR is located on the apical membrane of type A intercalated cells of the outer medullary collecting duct (OMCD) cells, where an increase in luminal [Ca^2+^] enhances H^+^ transport ^26, 29^. This effect is represented by setting α_OMCD,HATP_to be 2. Finally, CaSR is expressed on the apical membrane of inner medullary collecting duct (IMCD) cells, where an increase in luminal [Ca^2+^] reduces aquaporin 2 abundance. This effect is represented by setting α_IMCD,Pf_ to be -0.375.

### Transporter parameters

Model parameters that characterize Ca^2+^ transport is summarized in Table 1. Along the proximal tubule, Ca^2+^ is mainly reabsorbed via the paracellular pathway. Apical Ca^2+^ permeability is chosen so that the transcellular flux of Ca^2+^ accounts for ∼15% of total proximal tubule Ca^2+^ reabsorption. Along the thick ascending limb, Ca^2+^ reabsorption is entirely paracellular ^30^, driven by the favorable membrane potential established by K^+^ cycling into the lumen. Micropuncture experiments have indicated higher Ca^2+^ permeability along the cortical segment compared to the medullary segment ^31^.

**Table 1:** Calcium-specific parameters for all the segments along the superficial nephron. Values presented in (*) are adjusted and values presented in (**) denote that the parameter is the same for both male and female models. PT, proximal tubule; TAL, thick ascending limb; DCT2, late part of distal convoluted tubule; CNT, connecting tubule; CD, collecting duct; OMCD, outer medullary collecting duct; IMCD, inner medullary collecting duct.

| Parameter | Value |  |
| --- | --- | --- |
|  | Male | Female |
| **PT** |  |  |
| Permeability to $\text{Ca}^{2+}$ for Tight Junction | $20 \times 10^{-5} \text{ cm/s}^{76}$ | $100 \times 10^{-5} \text{ cm/s (*)}$ |
| Permeability to $\text{Ca}^{2+}$ for Basement Membrane | $60 \times 10^{-5} \text{ cm/s}^{77}$ | $60 \times 10^{-5} \text{ cm/s (**)}$ |
| Permeability to $\text{Ca}^{2+}$ for Apical Cell Membrane | $0.005 \times 10^{-5} \text{ cm/s (*)}$ | $0.03 \times 10^{-5} \text{ cm/s (*)}$ |
| PMCA maximum flux, $J_{\text{Ca}}^{\text{PMCA,max}}$ | $0.5 \times 10^{-9} (*)$<br>$\text{mmol/cm}^2/\text{s}$ | $2.5 \times 10^{-9} \text{ mmol/cm}^2/\text{s}$<br>$(*)$ |
| PMCA affinity to $\text{Ca}^{2+}$, $K_{\text{m,PMCA}}$ | $75.6 \text{ nM}^{78}$ | $75.6 \text{ nM (**)}$ |
| Reflection coefficient of tight junction to $\text{Ca}^{2+}$ | $0.89^{79}$ | $0.89 (**)$ |
| **TAL** |  |  |
| Maximum half concentration, $\text{EC}_{50}^{\text{n}}$ | $1.25 \text{ mM}^{27}$ | $1.25 (**)$ |
| Hill function coefficient, $n$ | $4^{80}$ | $4 (**)$ |
| Inhibitory coefficient, $\alpha_{\lambda}$ | $-4/7^{64}$ | $-4/7 (**)$ |
| Tight junction permeability in the absence of $\text{Ca}^{2+}$, $\lambda^*$ | $12 \times 10^{-5} \text{ cm/s}^{64}$ | $12 \times 10^{-5} \text{ cm/s (**)}$ |
| Inhibitory coefficient for NKCC2 activity, $\alpha_{\text{NKCC2}}$ | $-0.5^{81}$ | $-0.5 (**)$ |
| Inhibitory coefficient for potassium channel activity, $\alpha_{\text{ROMK}}$ | $-0.84^{29}$ | $-0.84 (**)$ |
| **DCT2-CNT** |  |  |
| PMCA maximum flux, $J_{\text{Ca}}^{\text{PMCA,max}}$ | $4.0 \times 10^{-9} \text{ mmol/cm}^2/\text{s}$ | $4.5 \times 10^{-9} \text{ mmol/cm}^2/\text{s}$<br>$(*)$ |
| PMCA affinity to $\text{Ca}^{2+}$, $K_{\text{m,PMCA}}$ | $42.6 \text{ nM}^{78}$ | $42.6 \text{ nM (**)}$ |
| Channel density, $N_{\text{TRPV5}}$ | DCT2: $20 \times 10^6$<br>$1/\text{cm}^2 (*)$,<br>CNT: $5 \times 10^6 / \text{cm}^2 (*)$ | DCT2: $35 \times 10^6 / \text{cm}^2$<br>Based on $^{34}$<br>CNT: $8.75 \times 10^6 / \text{cm}^2$<br>Based on $^{34}$ |
| Intracellular $[Ca^{2+}]$ for half-maximal block, $C_{inhib}^{TRPV5}$ | 74 nM <sup>78</sup> | 74 nM (**) |
| NCX1 maximum flux, $J_{Ca}^{NCX1,max}$ | $2,500 \times 10^{-9}$ mmol/cm<sup>2</sup>/s (*) | $2,750 \times 10^{-9}$ mmol/cm<sup>2</sup>/s (**) |
| NCX1 affinity to $Ca^{2+}$, $K_{m,NCX1}$ | 0.125 M <sup>82</sup> | 0.125 M (**) |
| Constant for voltage dependence, $\gamma$ | 0.35 | 0.35 (**) |
| Internal $Ca^{2+}$ half-saturation constant, $K_{m,Cai}$ | 3.59 M <sup>82</sup> | 3.59 M (**) |
| External $Ca^{2+}$ half-saturation constant, $K_{m,Cao}$ | 1.3 mM <sup>82</sup> | 1.3 mM (**) |
| Internal $Na^{+}$ half-saturation constant, $K_{m,Nai}$ | 12.29 mM <sup>82</sup> | 12.29 mM (**) |
| External $Na^{+}$ half-saturation constant, $K_{m,Nao}$ | 87.5 mM <sup>82</sup> | 87.5 mM (**) |
| Excitatory coefficient for NCC activity, $\alpha_{NCC}$ | 0.5 <sup>28</sup> | 0.5 (**) |
| **CD** |  |  |
| Tight Junction | $20 \times 10^{-5}$ cm/s <sup>83</sup> | $20 \times 10^{-5}$ cm/s (**) |
| Basement Membrane | $4.5 \times 10^{-5}$ cm/s | $4.5 \times 10^{-5}$ cm/s (**) |
| Promoting coefficient for apical HATPase activity of type A OMCD cells, $\alpha_{HATP}$ | +2 <sup>84</sup> | +2 (**) |
| Inhibitory coefficient for apical water permeability of IMCD cells, $\alpha_{pf}$ | -3/8 <sup>29</sup> | -3/8 (**) |

*Ex vivo* experiments using rabbit proximal tubule cells indicate that estradiol enhances Ca^2+^ reabsorption ^32^. Thus, transcellular and paracellular permeability and 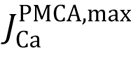 are assumed to be larger in the female model relative to male. No significant sex difference has been observed along the thick ascending limb in terms of Ca^2+^ transport ^33^. Along the distal tubular segments, female rats exhibit higher levels of both TRPV5 mRNA and protein expression compared to male rats, as reported in a study by Hsu et al.^34^ In the absence of sufficient sex-specific data, CaSR parameters are assumed the same for both sexes.

### Simulating NHE3 inhibition

In two separate simulations, expression of NHE3 along the proximal tubules was inhibited by 50% and 80%. In addition to NHE3, NHE2 has been shown to be expressed in the apical membrane of the cortical and medullary thick ascending limbs. Thus, in the 50% and 80% NHE3 inhibition simulations, we inhibited NHE expression along the thick ascending limbs by 25% and 40%, respectively. We assume that NHE3 was inhibited for a sufficiently long period for washout and changes in interstitial osmolarity to occur but before the manifestation of compensatory mechanisms. The baseline parameters correspond to a rat kidney in a mild antidiuretic state, such that urine osmolality is 771 mosmol/(kg H_2_O). When NHE3 is inhibited, fluid delivery to the loops of Henle and collecting ducts increases. The resulting significantly higher osmotic load presented to the concentrating mechanism is expected to overwhelm and impede the concentrating effect. Given these observations, we lowered the medullary interstitial concentrations of selected solutes in the NHE3 inhibition simulations; cortical interstitial concentration profiles were assumed to remain unaffected. For details, see Ref. ^35^.

### Simulating NKCC2 inhibition

NKCC2 is expressed on the apical membrane of the thick ascending limbs of the loops of Henle. We assumed that NKCC2 inhibition was administrated for long enough to significantly impair the kidney’s ability to generate an axial osmolality gradient. Thus, as in the NHE3 inhibition simulations, we lowered the interstitial fluid concentrations of selected solutes; for details, see Ref. ^36^. In addition to the thick ascending limb cells, NKCC2 is also expressed in the apical membrane of macula densa cells. Indeed, NKCC2-mediated transport constitutes the initial step in the signaling pathway between the macula densa cells and afferent arteriole smooth muscle cells. Targeted deletion of NKCC2 significantly attenuates the tubuloglomerular feedback (TGF) response ^37–39^. Thus, in the NKCC2 inhibition simulations, we assumed that SNGFR remained at baseline values, consistent with an experimental study in the rat ^40^.

### Simulating NCC inhibition

NCC is expressed on the apical membrane of the distal convoluted tubule. In the NCC inhibition simulations, baseline interstitial concentration profiles were used. Other model parameters remained at baseline values.

### Simulating ENaC inhibition

ENaC is expressed on the apical membrane of the last third of the distal convoluted tubules as well as the full length of the connecting tubules and collecting ducts. Administration of amiloride to rats has a sparing effect on Mg^2+^ reabsorption, raising intercellular [Mg^2+^], which has an inhibitory effect on TRPV5 ^41^. Since Mg^2+^ is not represented in the present models, we reduced TRVP5 expression by 10% in the case of 100% ENaC inhibition to reflect this effect in our model. As in the NCC inhibition simulations, baseline interstitial concentration profiles and non-ENaC parameters were used.

## Results

Using baseline sex-specific model parameters (Table 2), we compute steady-state solutions to the model equations. The models predict luminal fluid flow, hydrostatic pressure, luminal fluid solute concentrations, and, with the exception of the descending limb segment, cytosolic solute concentrations, membrane potential, and transcellular and paracellular water and solute fluxes, separately for the model superficial nephron of a female and male rat. Fig. 2 shows the delivery of key solutes [Na^+^, Ca^2+^, K^+^, Cl^−^, HCO_3_^−^, NH_4_^+^, urea, and titratable acid (TA)] and fluid to the inlets of individual nephron segments predicted by the female and male nephron models. The corresponding solute and fluid transport along individual nephron segments are shown in Fig. 3.

**Figure 2:**
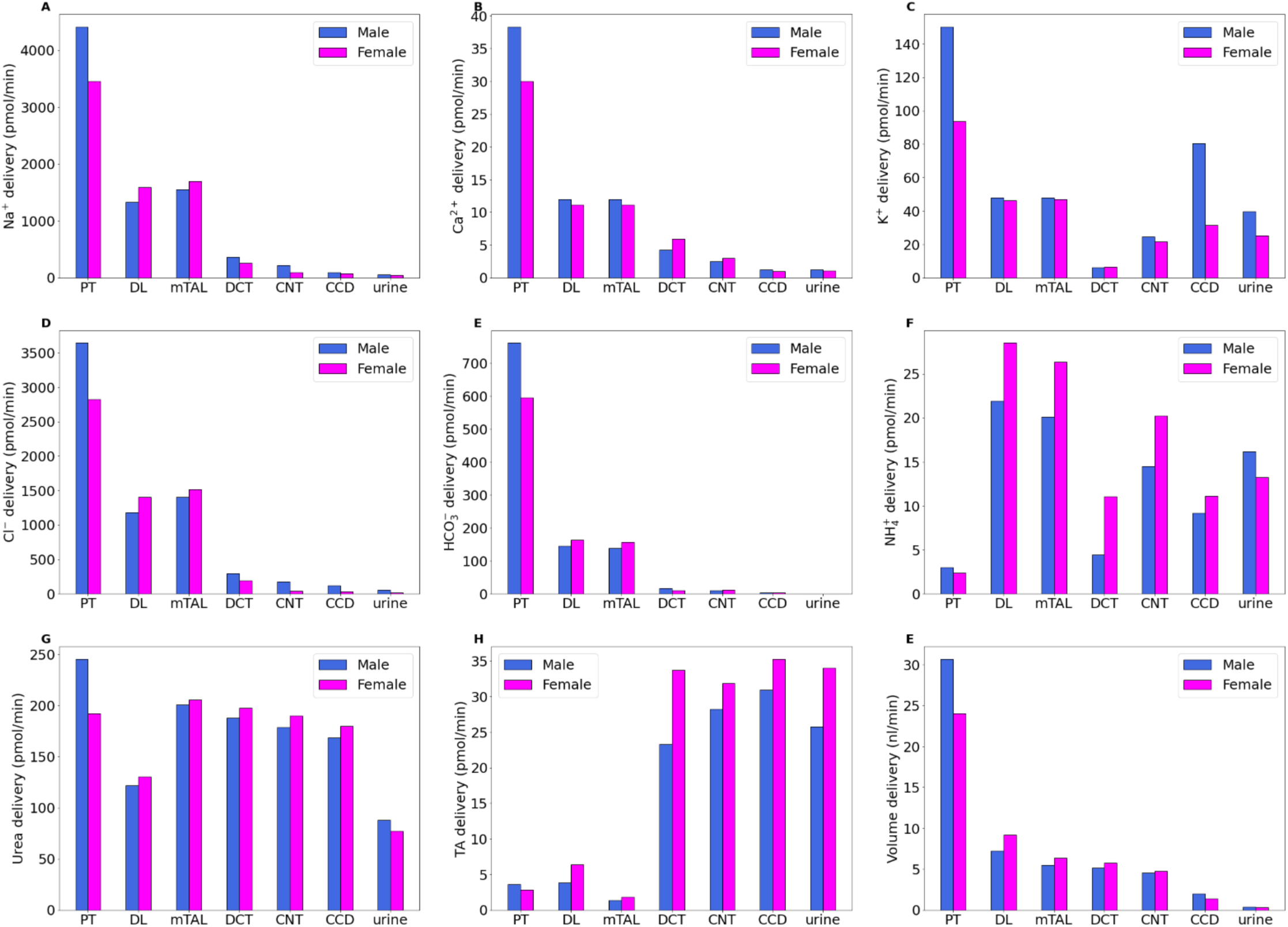
Delivery of key solutes (A–H) and fluid (G) to the beginning of individual nephron segments in male and female rats under baseline conditions. PT, proximal tubule; DL, descending limb; mTAL, medullary thick ascending limb; DCT, distal convoluted tubule; CNT, connecting tubule; CCD, cortical collecting duct; TA, titratable acid.

**Figure 3:**
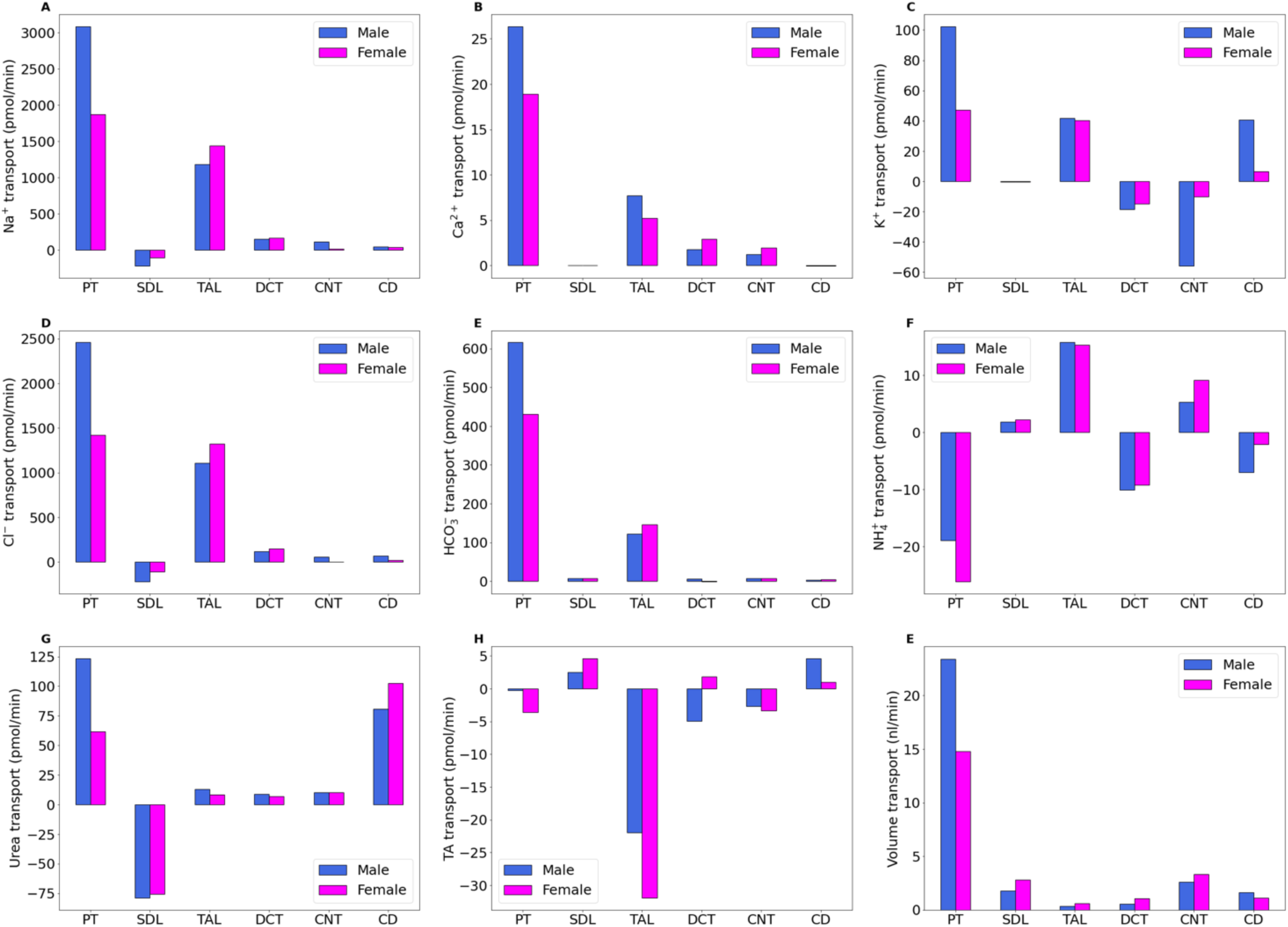
Net transport of key solutes (A–H) and fluid (E) along individual nephron segments, in male and female rats under baseline conditions. Transport is taken positive out of a nephron segment. PT, proximal tubule; DL, descending limb; TAL, thick ascending limb; DCT, distal convoluted tubule; CNT, connecting tubule; CD, collecting duct; TA, titratable acid.

**Table 2:** NaPi2, Na^+^−P_i_ cotransporter 2; P_Na_, Na^+^ permeability; P_Cl_, Cl^−^permeability; P_f_, water permeability; NKCC2, Na^+^−K^+^− Cl^−^cotransporter isoform 2; KCC, K^+^ − Cl^−^ cotransporter; ENaC, epithelial Na^+^ channel; P_K_, K^+^ permeability; PCT, proximal convoluted tubule; S3, proximal straight tubule; mTAL, medullary thick ascending limb; cTAL, cortical thick ascending limb; DCT, distal convoluted tubule; CNT, connecting tubule; CCD, cortical collecting duct; OMCD, outer medullary collecting duct; IMCD, inner medullary collecting duct; SNGFR, single nephron glomerular filtration rate.

| Parameter | Female-to-Male Ratio | Parameter | Female-to-Male Ratio |
| --- | --- | --- | --- |
| **PCT** |  | **S3** |  |
| NHE3 activity | 0.83 |  | 0.83 |
| NaPi2 | 0.75 |  | 0.75 |
| P<sub>Na</sub>, P<sub>Cl</sub> (paracellular) | 0.4 |  | 0.4 |
| P<sub>f</sub> (transcellular) | 0.64 |  | 0.64 |
| **mTAL** |  | **cTAL** |  |
| NKCC2 activity | 2 |  | 2 |
| KCC activity | 1.5 |  | 1.5 |
| Na<sup>+</sup>-K<sup>+</sup>-ATPase activity | 2 |  | 2 |
| Na<sup>+</sup> / H<sup>+</sup> exchanger | 0.8 |  | 0.8 |
| P<sub>Na</sub>, P<sub>Cl</sub> (paracellular) | 0.9 |  |  |
| **DCT** |  | **CNT** |  |
| NCC activity | 1.6 |  |  |
| Na<sup>+</sup>-K<sup>+</sup>-ATPase activity | 2 |  | 2 |
| Na<sup>+</sup>/H<sup>+</sup> exchanger | 0.85 |  | 0.9 |
| ENaC activity | 2 |  | 1.3 |
| P<sub>Na</sub>, P<sub>Cl</sub> (paracellular) | 1.4 |  | 1.4 |
| P<sub>f</sub> (transcellular) | 2 |  | 1.5 |
| **CCD** |  | **OMCD** |  |
| Na<sup>+</sup>-K<sup>+</sup>-ATPase activity | 1.5 |  | 1.1 |
| H<sup>+</sup>-K<sup>+</sup>-ATPase activity | 1 |  | 1.5 |
| Na<sup>+</sup>/H<sup>+</sup> exchanger | 0.9 |  | 0.9 |
| ENaC activity | 1.5 |  | 1.2 |
| P<sub>Na</sub>, P<sub>Cl</sub> (paracellular) | 1.4 |  | 1 |
| P<sub>K</sub> (apical) | 0.7 |  | 1.2 |
| P<sub>f</sub> (transcellular) | 2 |  | 2 |
| **IMCD** |  | **Morphology** |  |
| Na<sup>+</sup>-K<sup>+</sup>-ATPase activity | 1.2 | **PCT, S3** |  |
| H<sup>+</sup>-K<sup>+</sup>-ATPase activity | 1.5 | Length | 0.8 |
| P<sub>Na</sub>, P<sub>Cl</sub> (paracellular) | 0.5 | Diameter | 0.8 |
| P<sub>K</sub> (apical) | 2 | **Distal segments** |  |
| P<sub>Na</sub> (apical) | 0.5 | Length | 0.85 |
| P<sub>f</sub> (transcellular) | 2 | Diameter | 0.85 |
|  |  | Hemodynamics |  |
|  |  | SNGFR | 0.8 |

**Table 3:** Baseline and inhibition excretion values for Na^+^ and Ca^2+^. Transport and excretion values are given in pmol/min. Percentage changes from baseline values are shown in parentheses. Co-inhibition refers to the simultaneous inhibition of TRPV5, NCX1, and PMCA.

|  | PT transport |  | TAL transport |  | DCT transport |  | CNT transport |  | Urinary excretion |  |
| --- | --- | --- | --- | --- | --- | --- | --- | --- | --- | --- |
|  | Na<sup>+</sup> | Ca<sup>2+</sup> | Na<sup>+</sup> | Ca<sup>2+</sup> | Na<sup>+</sup> | Ca<sup>2+</sup> | Na<sup>+</sup> | Ca<sup>2+</sup> | Na<sup>+</sup> | Ca<sup>2+</sup> |
| **Base case** |  |  |  |  |  |  |  |  |  |  |
| **Male** | 3085 | 26 | 1180 | 7.7 | 154 | 1.8 | 117 | 1.3 | 53 | 1.2 |
| **Female** | 1869 | 19 | 1438 | 5.2 | 168 | 2.9 | 13 | 2.0 | 42 | 1.0 |
| **Ca<sup>2+</sup> Specific Inhibitions** |  |  |  |  |  |  |  |  |  |  |
| **Co-inhibition 100 (male)** | 3085 (0%) | 26 (0%) | 1180 (0%) | 7.7 (0%) | 153 (0%) | 0.0 (-99%) | 119 (+2%) | 0.0 (-99%) | 51 (-2%) | 4.6 (+269%) |
| **Co-inhibition 100 (female)** | 1869 (0%) | 19 (0%) | 1438 (0%) | 5.2 (0%) | 170 (+1%) | 0.0 (-99%) | 13 (+2%) | 0.0 (-99%) | 37 (-12%) | 6.1 (+499%) |
| **NCX1-100 (male)** | 3085 (0%) | 26 (0%) | 1180 (0%) | 7.7 (0%) | 154 (0%) | 1.7 (-8%) | 117 (0%) | 0.81 (-35%) | 52 (-1%) | 1.8 (+47%) |
| **NCX1-100 (female)** | 1869 (0%) | 19 (0%) | 1438 (0%) | 5.2 (0%) | 170 (+1%) | 1.7 (-40%) | 13 (0%) | 0.78 (-60%) | 39 (-7%) | 3.4 (+232%) |
| **PT-50 (male)** | 3085 (0%) | 24 (-7%) | 1180 (0%) | 9.4 (+22%) | 154 (0%) | 1.8 (0%) | 118 (+1%) | 1.3 (0%) | 54 (+2%) | 1.5 (+17%) |
| **PT-50 (female)** | 1869 (0%) | 19 (-1%) | 1437 (0%) | 5.0 (+5%) | 168 (0%) | 2.9 (0%) | 13 (0%) | 2.0 (0%) | 42 (0%) | 1.1 (+5%) |
| **TAL-50 (male)** | 3085 (0%) | 26 (0%) | 1182 (0%) | 5.8 (-25%) | 154 (0%) | 1.8 (+2%) | 117 (0%) | 1.3 (+4%) | 51 (-3%) | 3.0 (+146%) |
| **TAL-50 (female)** | 1869 (0%) | 19 (0%) | 1439 (0%) | 3.6 (-31%) | 168 (0%) | 3.0 (+1%) | 13 (-1%) | 2.0 (+4%) | 40 (-4%) | 2.5 (+144%) |
| **Na<sup>+</sup> Specific Inhibitions** |  |  |  |  |  |  |  |  |  |  |
| **NHE3-50 (male)** | 2529 (-18%) | 22 (-18%) | 1203 (+2%) | 7.8 (+1%) | 168 (+9%) | 1.7 (-4%) | 156 (+33%) | 1.2 (-5%) | 98 (+87%) | 2.2 (+78%) |
| **NHE3-50 (female)** | 1534 (-17%) | 16 (-16%) | 1312 (-8%) | 6.1 (+15%) | 220 (+31%) | 2.3 (-21%) | 66 (+410%) | 1.8 (-7%) | 69 (+66%) | 1.2 (+20%) |
| **NHE3-80 (male)** | 1708 (-44%) | 15 (-44%) | 1153 (-2%) | 7.3 (-4%) | 198 (+29%) | 1.8 (0%) | 194 (+66%) | 1.2 (-7%) | 177 (+236%) | 3.0 (+133%) |
| **NHE3-80 (female)** | 1091 (-41%) | 12 (-37%) | 1265 (-12%) | 5.9 (+13%) | 247 (+47%) | 3.0 (+1%) | 154 (+1086%) | 1.8 (-6%) | 196 (+370%) | 2.3 (+126%) |
| **NKCC2-70 (male)** | 3066 (0%) | 26 (0%) | 961 (-18%) | 6.1 (-21%) | 163 (+6%) | 1.7 (-7%) | 154 (+32%) | 1.2 (-3%) | 120 (+128%) | 3.2 (+156%) |
| **NKCC2-70 (female)** | 1808 (-3%) | 18 (-4%) | 1267 (-11%) | 4.2 (-20%) | 204 (+21%) | 2.7 (-9%) | 63 (+384%) | 1.9 (-2%) | 103 (+148%) | 3.3 (+220%) |
| **NKCC2-100 (male)** | 3049 (-1%) | 26.1 (-1%) | 240 (-79%) | -1.0 (backleak) | 184 (+19%) | 1.4 (-19%) | 229 (+95%) | 0.89 (-29%) | 609 (+1057%) | 11.0 (+785%) |
| **NKCC2-100 (female)** | 1774 (-5%) | 17 (-8%) | 287 (-80%) | -2.9 (backleak) | 241 (+43%) | 2.2 (-26%) | 204 (+1469%) | 1.5 (-22%) | 838 (+1914%) | 12 (+1068%) |
| **NCC-70 (male)** | 3085 (0%) | 26 (0%) | 1180 (0%) | 7.7 (0%) | 107 (-30%) | 1.9 (+3%) | 145 (+23%) | 1.1 (-10%) | 64 (+21%) | 1.3 (+5%) |
| **NCC-70 (female)** | 1845 (-1%) | 19 (-1%) | 1446 (0%) | 5.2 (0%) | 130 (-22%) | 2.7 (-8%) | 45 (+247%) | 1.8 (-6%) | 53 (+27%) | 1.4 (+37%) |
| **NCC-100 (male)** | 3085 (0%) | 26 (0%) | 1180 (0%) | 7.7 (0%) | 49 (-68%) | 1.8 (-2%) | 172 (+46%) | 0.98 (-21%) | 83 (+58%) | 1.6 (+25%) |
| **NCC-100 (female)** | 1845 (0%) | 19 (0%) | 1446 (0%) | 5.2 (0%) | 31 (-81%) | 2.0 (-30%) | 106 (+716%) | 1.6 (-18%) | 79 (+90%) | 2.3 (+123%) |
| **ENaC-70 (male)** | 3085 (0%) | 26 (0%) | 1180 (0%) | 7.7 (0%) | 150 (-2%) | 2.3 (+28%) | 54 (-53%) | 1.6 (+30%) | 93 (+76%) | 0.35 (-71%) |
| **ENaC-70 (female)** | 1869 (0%) | 19 (0%) | 1446 (0%) | 5.2 (0%) | 171 (+1%) | 3.5 (+19%) | -7.8 (backleak) | 2.1 (+8%) | 69 (+66%) | 0.28 (-73%) |
| **ENaC-100 (male)** | 3085 (0%) | 26 (0%) | 1180 (0%) | 7.7 (0%) | 147 (-4%) | 2.4 (+34%) | -24 (backleak) | 1.8 (+46%) | 173 (+228%) | 0.02 (-98%) |
| **ENaC-100 (female)** | 1869 (0%) | 19 (0%) | 1446 (0%) | 5.2 (0%) | 168 (0%) | 3.8 (+26%) | -37 (backleak) | 2.2 (+11%) | 115 (+176%) | 0.01 (-99%) |

### Proximal tubule transport

The models predict that proximal tubules of male and female nephrons reabsorb 70% and 55%, respectively, of filtered Na^+^, primarily via NHE3 (apical) and Na^+^-K^+^-ATPase (basolateral). The fractional reabsorption values are substantially lower in female nephrons because of their smaller transport area and lower NHE3 activity. The high water-permeability of the proximal tubule epithelium facilitates water removal from the luminal fluid following Na^+^ reabsorption. Significant water and Na^+^ reabsorption also occurs through the paracellular pathway ^42, 43^. Overall, the proximal tubules of male and female nephrons reabsorb 76% and 62% of filtered volume.

The sex difference in Na^+^ transport carries over to Ca^2+^ transport, inasmuch as the majority of the filtered Ca^2+^ is reabsorbed by the proximal tubule via a passive paracellular process, driven by Na^+^ and subsequent water reabsorption, which establishes a favorable electrochemical gradient for Ca^2+^ reabsorption. Ca^2+^ reabsorption is mediated by Cldn-2 and Cldn-12, and by solvent drag. The models predict that 68% and 61%, respectively, of the filtered Ca^2+^ is reabsorbed along the proximal tubule in the male and female rat nephron, in parallel to the fractional Na^+^ reabsorption (see above). Luminal fluid Ca^2+^ concentration in the proximal convoluted tubule is predicted to remain close to plasma, consistent with micropuncture findings ^44, 45^.

### Thick ascending limb transport

The majority of the Na^+^ exiting the proximal tubule is reabsorbed along the thick ascending limb. Specifically, fractional Na^+^ reabsorption is predicted to be 27% and 42% in the male and female thick ascending limb, respectively; this corresponds to a 22% higher net Na^+^ transport in the female thick ascending limb, despite its smaller transport area, because of the assumption of higher NKCC2, KCC, and Na^+^-K^+^-ATPase activities in female.

Calcium reabsorption along the thick ascending limb is entirely paracellular, driven by the positive transepithelial potential established in large part by the apical recycling of K^+^. Unlike Na^+^, less Ca^2+^ is reabsorbed along the medullary versus cortical thick ascending limbs. This is because the higher interstitial [Ca^2+^] in the outer medulla, especially in the inner stripe, causes the CaSR to lower tight-junction Ca^2+^ permeability in the medullary segment. Consequently, the medullary and cortical thick ascending limb segments of the male rat nephron reabsorb 8% and 14% of the filtered Ca^2+^. Despite the higher net and fractional Na^+^ reabsorption in female, its medullary and cortical thick ascending limb segments reabsorb less Ca^2+^ compared to male: 6% and 12% of the filtered load, respectively. This difference may be attributed to the lower intracellular [K^+^] in females, resulting in lower apical K^+^ secretion and a less favorable membrane potential to drive Ca^2+^ reabsorption.

### Distal segments

In contrast to the upstream segments, Ca^2+^ transport along the distal convoluted tubule and connecting tubule occurs through the transcellular pathway: Ca^2+^ first passes the apical membrane through TRPV5 and is then extruded from the basolateral membrane primarily by the NCX1, with the rest by the PMCA. In the male model, the distal convoluted tubule and connecting tubule are predicted to mediate 4.7% and 3.2% of the filtered Ca^2+^, respectively. In the female model, TRPV5 abundance is assumed to be considerably higher ^34^(see Table 1), with its activity further enhanced due to the lower luminal fluid pH compared to male. As such, considerably more Ca^2+^, amounting to 9.7% and 6.5% of its filtered load, is reabsorbed by the distal convoluted tubule and connecting tubule in the female model. Ca^2+^ transport is negligible along the collecting duct.

Taken together, the models predict a urine output of 0.38 and 0.31 nl/min for the male and female rat nephron, respectively, and urinary Na^+^ excretion of 52.6 and 41.6 pmol/min, and urinary excretion of Ca^2+^ of 1.24 and 1.02 pmol/min. These results are summarized in Table 3.

### Co-inhibition of calcium transport along late distal convoluted tubule and connecting tuble

Complete inhibition of TRPV5 expression along with NCX1 and PMCA eliminates Ca^2+^ reabsorption along the distal convoluted tubule and connecting tubule and substantially increases Ca^2+^ delivery to the collecting duct, resulting in hypercalciuria. As shown in Table 3, urinary Ca^2+^ excretion is predicted to increase more than two-fold (269%) in the male model, and even more (by 499%) in female due to the higher TRPV5 activity (Table 3). Inhibition of TRPV5 renders ΔV_te_ more lumen-positive and enhances Na^+^ reabsorption, decreasing urinary Na^+^ excretion, although that effect is small compared to changes in Ca^2+^ excretion (by 5% and 12% in male and female, respectively).

### NCX1 Inhibition

Ca^2+^ extrusion from the late distal convoluted tubule and connecting tubule cells is mediated primarily through the NCX1 with the remainder via the PMCA (∼70% for both male and female models). Thus, complete inhibition of NCX1 substantially decreases Ca^2+^ reabsorption along these segments (by 22% and 48% in the male and female models, respectively), as PMCA is unable to fully compensate. Eventually, this leads to major increases in urinary Ca^2+^ reabsorption, in male by 47% and in female, which has higher NCX1 activity, by 232%. As in the case of co-inhibition, NCX1 inhibition promotes Na^+^ reabsorption and increases urinary Na^+^ excretion (Table 3).

### Decreasing proximal tubule and thick ascending limb Ca^2+^ transport

Simulations were conducted to investigate the effects of decreasing proximal tubule paracellular Ca^2+^ permeability by 50%. Transport parameters were kept at baseline values in other segments. The models predict that the thick ascending limbs compensate by elevating its Ca^2+^ transport, resulting in significant but not drastic increases in urinary Ca^2+^ excretion. In the male model, by decreasing the paracellular permeability of calcium in the proximal tubule, it leads to a greater increase in calcium excretion due to lower reabsorption in this segment compared to the female model (17% in male and 5% in female).

When thick ascending limb paracellular Ca^2+^ permeability was decreased by 50%, the downstream late distal convoluted tubules and connecting tubules were unable to sufficiently compensate for the reduction in Ca^2+^ transport. Consequently, the urinary Ca^2+^ excretion is predicted to increase by ∼140% in both male and female models. When either proximal tubule or thick ascending limb Ca^2+^ transport was partially inhibited, the effects on Na^+^ transport and excretion are negligible; see Table 3.

### NHE3 inhibition

Given that the proximal tubule Ca^2+^ transport is driven, directly and indirectly, by the NHE3-mediated Na^+^ reabsorption, we conduct separate simulations to assess the impacts of 50% and 80% inhibition of NHE3. The resulting segmental delivery and transport of Ca^2+^ and Na^+^ are shown in Figs. 4A-4D and segmental transport values are summarized in Table 3 for both male and female.

**Figure 4:**
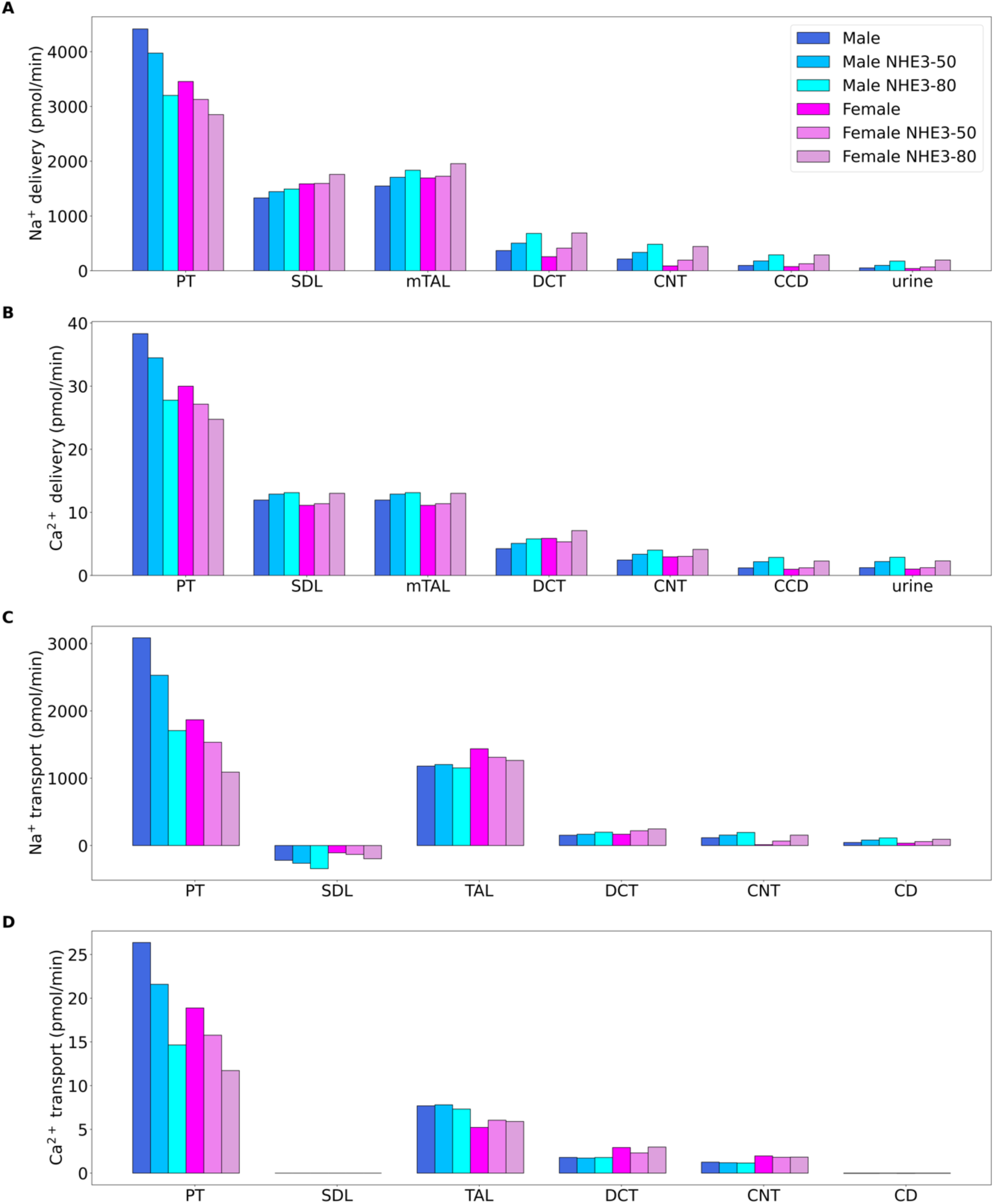
Comparison of segmental solute delivery of (A) Na^+^ and (B) Ca^2+^ and segmental solute transport of (C) Na^+^ and (D) Ca^2+^ in male and female, obtained for base case, and for 50% decrease in NHE3 activity (NHE3-50) and 80% decrease in NHE3 activity (NHE3-80); PT, proximal tubule; SDL, short descending limb; mTAL, medullary thick ascending limb; DCT, distal convoluted tubule; CNT, connecting tubule; CD, collecting duct; CCD, cortical collecting duct.

The baseline models predict that NHE3 mediates 87% and 81% of proximal tubule Na^+^ transport in male and female models, respectively. The effects of inhibiting NHE3 in the rat kidneys have been described in Ref. ^46^. When NHE3 is inhibited by 50%, the model proximal tubule reabsorbs 18% and 15% less Na^+^ in male and female, respectively. Because Ca^2+^ reabsorption is driven by that of Na^+^, this results in similar reductions in proximal tubule Ca^2+^ transport in both sexes. Qualitatively similar effects, but larger percentages, are predicted when NHE3 is inhibited by 80%.

With NHE3 inhibition, Na^+^ delivery to the thick ascending limbs is slightly elevated, despite a reduction in SNGFR. Taken in isolation, the higher delivery would enhance segmental transport. However, the NHE activity along the thick ascending limb is partially reduced. In male, these two competing factors result in essentially no change in thick ascending limb Na^+^ transport, but a larger fraction of which is now mediated by NKCC2. This in turn results in a small increase in K^+^ recycling and thus a more favorable membrane potential for Ca^2+^ transport. Consequently, with NHE3 inhibition, the thick ascending limb in the male model is predicted to reabsorb 5% more Ca^2+^, although the increase in Ca^2+^ transport becomes negligible at 80% inhibition. In the female model, the effect of the reduced NHE activity dominates and thick ascending limb Na^+^ transport is predicted to be significantly reduced. However, with the higher NKCC2 activity in female, K^+^ recycling and membrane potential increase more than in male; as such, Ca^2+^ transport along the thick ascending limb is predicted to increase by 15% and 7%, respectively, at 50% and 80% NHE3 inhibition.

Taken together, these segmental responses result in major natriuretic effects. Similarly, urinary Ca^2+^ excretion is predicted to increase substantially, to more than twice the baseline value when NHE3 is inhibited by 80%; see Table 3.

### NKCC2 inhibition

Next, we simulate 70% and 100% inhibition of NKCC2, which mediates Na^+^ transport along the thick ascending limbs, and indirectly drives paracellular Ca^2+^ reabsorption. Under baseline conditions, NKCC2 was predicted to mediate 88% of renal Na^+^ transport in the female nephron compared with 83% in the male nephron.

Inhibition of NKCC2 has limited direct impact on upstream proximal tubular transport, as we assume that TGF is not activated. Thick ascending limb Na^+^ transport fell similarly in both sexes when NKCC2 was inhibited: by 18% and 12% in male and female rats when NKCC2 was inhibited by 70% and by ∼80% in both sexes when NKCC2 was inhibited by 100% (Fig. 5C).

**Figure 5:**
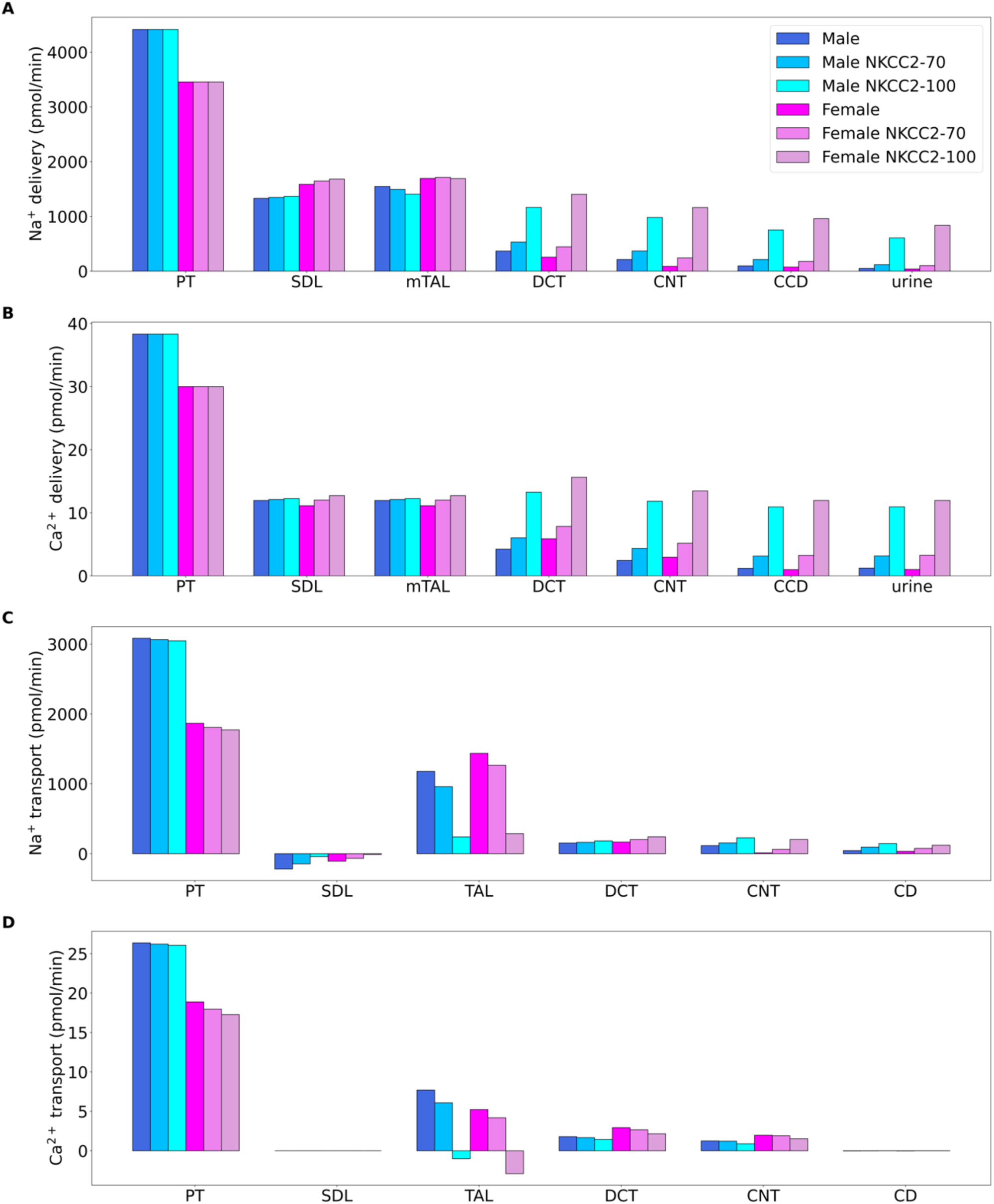
Comparison of segmental solute delivery of (A) Na^+^ and (B) Ca^2+^ and segmental solute transport of (C) Na^+^ and (D) Ca^2+^ in male and female, obtained for base case, and for 70% decrease in NKCC2 activity (NKCC2-70) and 100% decrease in NKCC2 activity (NKCC2-100); PT, proximal tubule; SDL, short descending limb; mTAL, medullary thick ascending limb; DCT, distal convoluted tubule; CNT, connecting tubule; CD, collecting duct; CCD, cortical collecting duct;

In addition, substantially less K^+^ is reabsorbed along the thick ascending limbs in both sexes. In particular, inhibition of NKCC2 decreases K^+^ exit through the apical membrane, thereby reducing the driving force for Ca^2+^ reabsorption. With 70% NKCC2 inhibition, thick ascending limb Ca^2+^ transport decreases by 16% and 23% in male and female, respectively; with complete inhibition, there is a back leak of Ca^2+^ (Fig. 5D).

NKCC2 inhibition increases Na^+^ delivery and transport along the distal segments (Figs. 5A and 5C). Taken in isolation, that would promote Ca^2+^ reabsorption. However, when NKCC2 is inhibited, thick ascending limb NHE transport increases, lowering luminal fluid pH and TRPV5 conductance. As a result, Ca^2+^ transport along the distal convoluted tubule and connecting tubule is predicted to decrease (Table 3). Taken together, NKCC2 inhibition leads to multifold increases in urinary excretion of Na^+^ and Ca^2+^ in both sexes (Table 3).

### NCC Inhibition

In the next set of simulations, we examined the effects of inhibiting NCC by 70% and 100% on tubular transport. NCC is present at the apical membrane of distal convoluted tubule, where along the last third of the segment (DCT2) it is co-localized with TRPV5. The predicted segmental Na^+^ and Ca^2+^ delivery and transport are shown in Figs. 6A-6D. NCC inhibition has no effect on segments upstream of the distal convoluted tubule. NCC inhibition considerably reduces Na^+^ transport along the distal convoluted tubule. The resulting higher luminal [Na^+^] increases the driving force for other apical Na^+^ transporters, notably the NHE2. The model predicts that with 70% and 100% NCC inhibition NHE2-mediated Na^+^ transport increases by 17% and 24% in the male model; similar increases are predicted for female. The consequent acidification of the luminal fluid inhibits TRPV5. These two competing effects result in minor changes in Ca^2+^ transport along the distal convoluted tubule in male. Because TRPV5 abundance is higher in female, NCC inhibition results in larger reductions in Ca^2+^ transport along that same segment compared to male (Fig. 6D, Table 3). Following NCC inhibition, the higher Na^+^ delivery to the connecting tubule increases Na^+^ reabsorption via ENaC. This results in a less favorable voltage gradient for transcellular Ca^2+^ reabsorption, affecting both sexes similarly. Inhibition of NCC by 70% and 100% results in an overall increase in urinary Ca^2+^ excretion by 5% and 25%, respectively, in male, and by 37% and 123%, respectively, in female. The larger impact of NCC inhibition on Ca^2+^ excretion in female is due to their higher baseline fractional reabsorption of Ca^2+^ by the DCT2 and the stronger TRPV5 inhibition.

**Figure 6:**
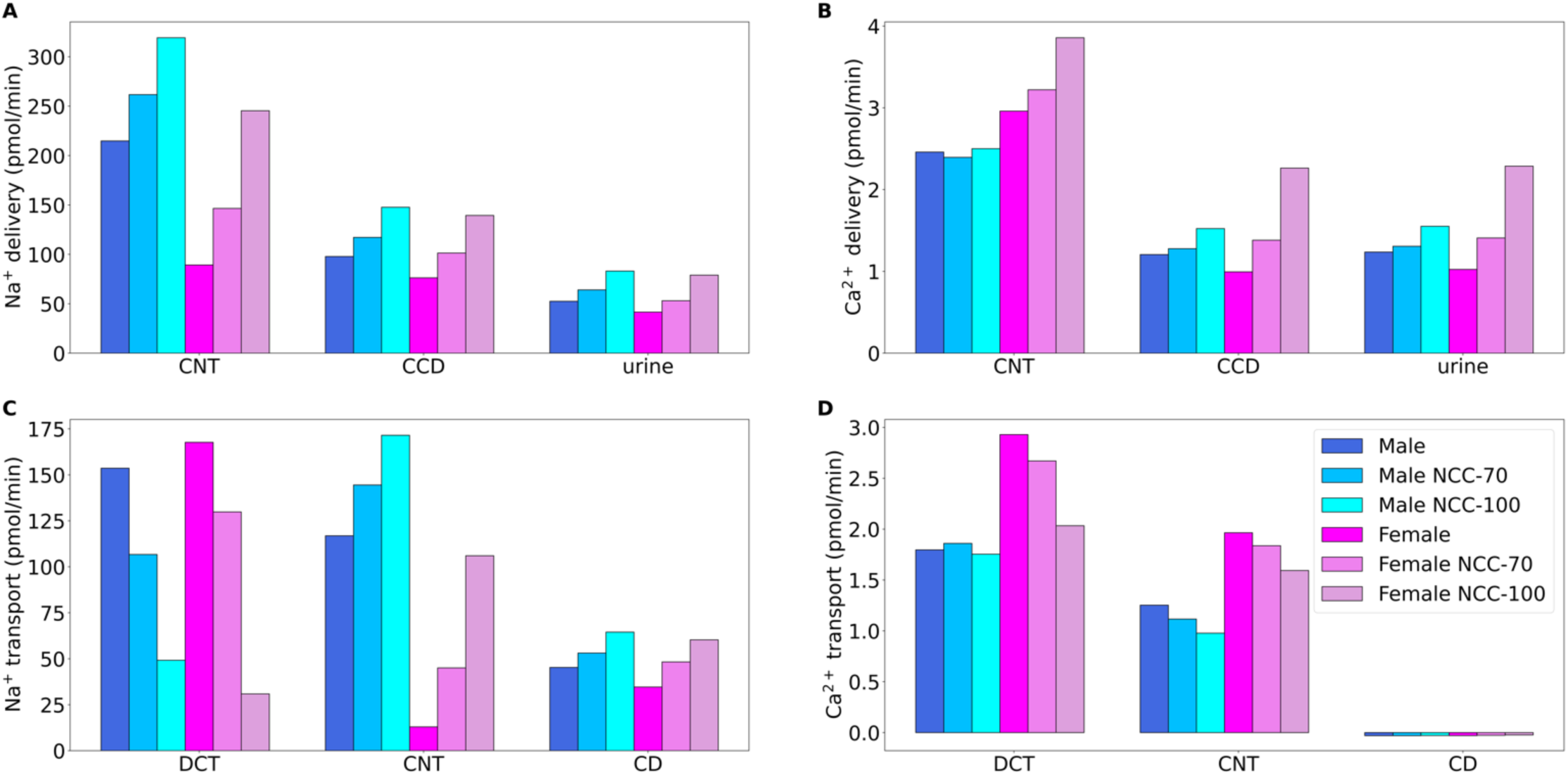
Comparison of segmental solute delivery of (A) Na^+^ and (B) Ca^2+^ and of segmental solute transport of (C) Na^+^ and (D) Ca^2+^ in male and female, obtained for base case, and for 70% decrease in NCC activity (NCC-70) and 100% decrease in NCC activity (NCC-100); CNT, connecting tubule; CD, collecting duct; CCD, cortical collecting duct.

### ENaC inhibition

ENaC is expressed on the apical membrane of the DCT2, connecting tubules and collecting ducts. Given the coupling between Ca^2+^ reabsorption and ENaC-mediated Na^+^ transport along these segments, we conducted separate simulations in which ENaC is inhibited by 70% and 100%. Recall also that TRPV5 activity is reduced to account for the effect of Mg^2+^. Segmental delivery and transport are summarized in Figs. 7A-7B and 7C-7D, along with transport values summarized in Table 3. ENaC inhibition hyperpolarizes the apical membrane, promoting Ca^2+^ reabsorption. As a result, Ca^2+^ excretion drops substantially by 71% in male and by 73% in female when ENaC is inhibited by 70%, and to almost zero with complete inhibition. The predicted reduction in Ca^2+^ excretion with 70% ENaC inhibition is consistent with observations in male rat experiments ^47^.

**Figure 7:**
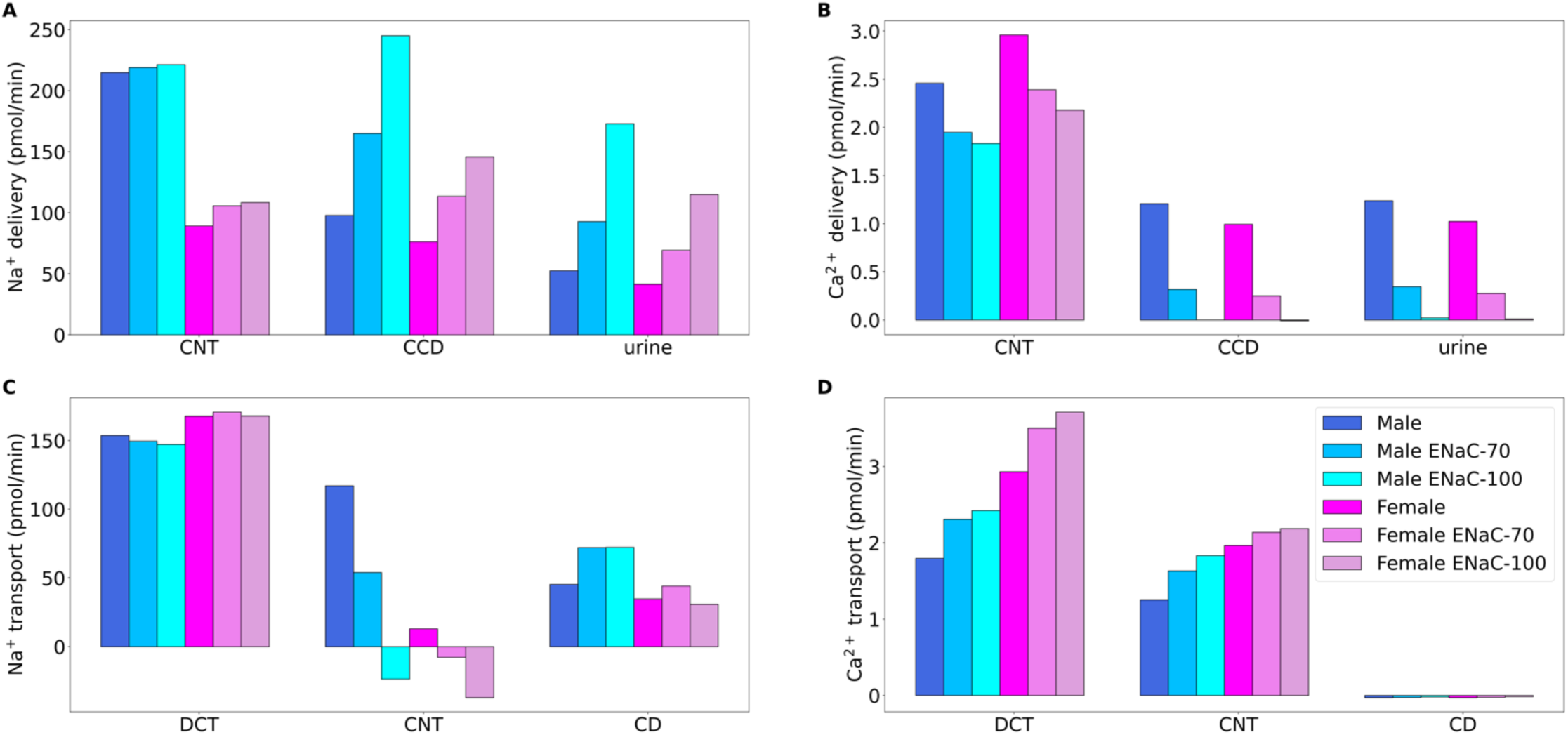
Comparison of segmental solute delivery of (A) Na^+^ and (B) Ca^2+^ and of segmental solute transport of (C) Na^+^ and (D) Ca^2+^ for male and female models, obtained for base case, and for 70% decrease in ENaC activity (ENaC-70) and 100% decrease in ENaC activity (ENaC-100); CNT, connecting tubule; CD, collecting duct; CCD, cortical collecting duct.

## Discussion

Luminal [Ca^2+^] in the accessible proximal convoluted tubule was measured to be close to that in the ultrafiltrate ^45^, evidence that the majority of proximal tubule Ca^2+^ reabsorption occurs via a passive paracellular process, driven by Na^+^ reabsorption mediated primarily by the NHE3, and by subsequent water reabsorption. The present models assume that the parameters that characterize Ca^2+^ transport along the proximal tubule have lower values in male compared to female (Table 1). Furthermore, sex differences in other transporters are incorporated. In particular, NHE3 activity is assumed to be significantly lower in female ^7, 46, 48^. Consequently, model simulations predict that the majority (68%) of the filtered Ca^2+^ is reabsorbed along the male proximal tubule, but fractional Ca^2+^ reabsorption is substantially lower (61%) in female.

The inhibition of calcium transporters across both the apical and basolateral membranes results in a notable almost 3-fold increase in calcium excretions in male and 5-fold increase female models. However, when focusing solely on the inhibition of NCX1, increased calcium excretion is particularly significant, amounting to a 47% and 232% increase in male and female models. Thus, emphasizing the important role of NCX1 in the control of calcium excretion.

Given that NHE3 is the main driver of proximal tubule Ca^2+^ reabsorption, we investigated the effect of NHE3 inhibition on renal Ca^2+^ transport. Model simulations indicate that NHE3 inhibition would lead to significant increases in urinary Ca^2+^ excretion (Table 3), consistent with the calcinuria reported in *Nhe3*-deficient mice ^49^. The effect was predicted to be smaller in female due to its relatively lower NHE3-mediated Na^+^ transport.

A common treatment for hypertension is diuretics, which increase urine output by altering the Na^+^ transport function of specific tubular segments. We investigated the effect on renal Ca^2+^ transport of three classes of diuretic medications: loop, thiazide, and K^+^-sparing diuretics. Loop diuretics inhibit NKCC2 on the apical membrane of the thick ascending limb by competing for the Cl^−^ binding site. NKCC2 is the primary mechanism that generates the lumen-positive transepithelial voltage gradient across the thick ascending limb epithelium. Thus, model simulations predict that the administration of loop diuretics diminishes this voltage gradient and significantly attenuates Ca^2+^ transport along the thick ascending limb. This result is consistent with the findings that administration of loop diuretics such as furosemide promotes hypercalciuria in experimental animals ^50–52^ and in human subjects ^53^.

Thiazide diuretics inhibit NCC on the distal convoluted tubule. By inducing NaCl loss, thiazides lower blood pressure and is often recommended as a first-line treatment for hypertensive patients. Given the coupling of Na^+^ and Ca^2+^ transport, model simulations predict Ca^2+^ loss following NCC inhibition (Table 3), consistent with the response of healthy subjects in the first 12 hours following hydrochlorothiazide administration ^53^. However, urinary Ca^2+^ excretion fell below baseline in the 12-24 hour period ^53^. Similar to the latter response, thiazides have been reported to promote hypocalciuria and may lead to hypercalcemia in some patients ^54, 55^. Hence, thiazides are often prescribed to alleviate hypocalcemia by increasing renal Ca^2+^ and to treat kidney stone disease by reducing urinary Ca^2+^ ^56, 57^. Similar effects were reported in rodents following chronic thiazide treatment and in mouse models of Gitelman syndrome ^58–60^. The mechanisms that lead to hypocalciuria following longer-term thiazide treatment or in Gitelman patients may be attributable to alterations in proximal transport mechanisms ^61^. Thiazides administration causes natriuresis and diuresis. The resulting volume contraction enhances proximal tubular Na^+^ reabsorption by pressure natriuresis. Because Ca^2+^ transport in the proximal tubule parallels that of Na^+^, enhanced Na^+^ transport leads to Ca^2+^ hyperabsorption and eventually hypocalciuria. Model simulations that represent pressure natriuresis by including elevated NHE3 activity in addition to NCC inhibition predict a reduction in urinary Ca^2+^ excretion (results not shown).

K^+^-sparing diuretics inhibit ENaC on the connecting tubule and collecting duct. Administration of amiloride to rats inhibited both Na^+^ and fluid reabsorption from the late distal convoluted tubule, but also augmented Ca^2+^ reabsorption, resulting in a reduced fractional excretion of Ca^2+^ ^62^. Following an administration of triamterene to healthy subjects, an initial Ca^2+^ loss was reported, but that changed after 6 h to hypocalciuria ^63^. This time-dependent response is similar to that observed in thiazide diuretics and may involve alternations in proximal transport (see above). Additionally, as noted in METHODS, administration of amiloride to rats increases Mg^2+^ reabsorption and intercellular [Mg^2+^], inducing an inhibitory effect on TRPV5 and attenuating Ca^2+^ reabsorption ^41^. When partial TRPV5 inhibition was included in the model’s ENaC inhibition simulations, hypocalciuria was predicted (Table 3). In contrast, without the TRPV5 inhibitory effect, the models would predict hypercalciuria (results not shown).

Recent computational models of kidney function have advanced our understanding of renal tubular function by accounting for solute and water transport, revealing insights into transport pathways, driving forces, and coupling mechanisms. That said, a challenge in the development of the present models is that some transport pathways are not yet fully characterized. For instance, the models represent the NCX1.1 isoform, which is the predominant isoform in cardiac cells and has a known kinetic model, instead of including a kinetic model for the NCX1.3 isoform, which is the predominant isoform in renal epithelial cells but is much less well characterized. Moreover, incorporating newly discovered proteins in specific nephron segments can be challenging when their functional importance is unclear, such as in the case of CaSR. The models represent the effects of CaSR along various segments, including the thick ascending limb, distal convoluted tubule, and the collecting duct. However, how CaSR may regulate proximal tubule cell transport has yet to be fully understood ^64^.

The present models simulate electrolyte and water transport along a superficial nephron. Superficial nephrons account for only 2/3 of the nephron population in a rat kidney; the remainder are made up of juxtamedullary nephrons whose loops of Henle reach into differing depths of the inner medulla. Single-nephron glomerular filtration rate, transport area, and transporter activities differ between the two classes of nephrons. The present study considers a superficial nephron model to understand segmental transport more clearly. In future studies, a kidney model that represents both types of nephrons ^65–68^ can more accurately predict urinary excretion rates.

Despite the limitations, the present models provide an essential component in understanding Ca^2+^ balance under physiological and pathophysiological conditions. The female model can be modified to simulate kidney function in a pregnant or lactating rat ^69^. During pregnancy, there are significant changes in hormone levels, along with structural changes such as hyperfiltration, tubular hypertrophy, and alternations in transporter abundance in the kidney which can affect renal Ca^2+^ handling. For example, the hormone estrogen can increase the activity of Ca^2+^ transporters in the kidneys, leading to increased Ca^2+^ reabsorption ^32^. On the other hand, the hormone progesterone can decrease Ca^2+^ reabsorption by promoting Ca^2+^ excretion ^70^. Moreover, during lactation, there is an increased demand for Ca^2+^ to support milk production ^71^, which can further affect renal Ca^2+^ handling. Thus, further investigation is needed to better understand the mechanisms underlying pregnancy- and lactation-related changes in renal Ca^2+^ handling, and the current model can be extended to explore these effects.

The effects of diabetes on renal Ca^2+^ handling have been observed in clinical trials ^72^ and animal experiments ^73^. In diabetes the kidney undergoes several changes such as hyperfiltration, tubular hypertrophy, and alterations in renal transporter abundance as previously investigated in renal epithelial solute transport models ^74, 75^, resulting in increased Na^+^ excretion. As Ca^2+^ transport is closely linked to Na^+^ transport, alterations in Na^+^ handling in diabetes impact renal Ca^2+^ handling as well. Moreover, studies in streptozotocin-induced diabetic rats have shown changes in protein expression, such as increased expression of TRPV5 ^73^. Therefore, investigating renal Ca^2+^ handling in male and female rats with diabetes could provide further insights into the complex interactions between diabetes and renal Ca^2+^ handling.

